# Compression Detects Changes in Spiking Neural Data from Cortical Lesions

**DOI:** 10.64898/2026.01.27.701628

**Authors:** Alice Tor, Yuxin Wu, Stephen E Clarke, Lisa Yamada, Tsachy Weissman, Paul Nuyujukian, the Brain Interfacing Laboratory

## Abstract

1

**Objective:** The complexity of neural data changes as the brain processes information during events. Universal lossless compression algorithms, which are broadly applicable and grounded in information theory, identify and exploit redundancies in data in order to compress it to essentially-optimal sizes regardless of underlying statistics. These algorithms may be used to conveniently and efficiently estimate a given signal’s Shannon entropy rate, a biologically relevant measure of the complexity of a signal. It is therefore natural to explore their effectiveness in the analysis of spiking neural data.

**Approach:** This work focuses on using compression to analyze recordings (96-channel Utah arrays) taken from motor cortex of animals performing reaching tasks for three days before and three days after administering electrolytic lesions (Subject U: 4 lesions, H: 3). In particular, we use the inverse compression ratio (ICR), which compares the sizes of compressed and uncompressed data to estimate the amount of statistically unique information. We calculate ICR with temporally-independent lossless compression (gzip) and temporally-dependent lossy compression (H.264, MPEG-2). Compression-based ICR was compared to single-neuron measures used to understand spiking data, such as average firing rates and Fano factor. Compression is also compared to common dimensionality reduction techniques, principal component analysis (PCA) and factor analysis (FA).

**Main Results:** Statistical tests on aggregate data comparing each metric before and after lesioning reveal that ICR is able to significantly (Mann-Whitney U test, p *<* 0.01) detect lesions with higher accuracy than single-neuron metrics, but not dimensionality reduction (ICR methods: 85.7%, single-neuron methods: 78.6%, dimensionality reduction: 100%). Additionally, statistical results on the same data show that ICR metrics remain more stable than single-neuron methods after lesion. The bitrate parameter of lossy compression algorithms is swept to better understand the effect of information rates and “optimal” compression on lesion detection performance. Our conclusions are confirmed by the same analyses performed on several different simulated neural datasets.

**Significance:** These results suggest that compression algorithms may be a useful tool to detect and better understand perturbations to the underlying structure of neural data. Information-theoretic analyses may complement techniques like dimensionality reduction and firing rate tuning as a convenient and useful tool to characterize neural data.

## 2 Introduction

Broad technological advances over the past few decades have led to the rapid development of many in-vivo neural recording devices. The number of neurons neuroscientists have access to continues to increase exponentially, drawing comparisons to a Moore’s law for neuroscience [1]. In particular, the advancement of implanted electrode arrays has dramatically shifted the field of systems neuroscience from theories based on individually recorded neurons to those based on the systems-level activity of populations of neurons [2–4]. The translation of high-dimensional neural activity to relatively simple tasks implies that there is significant redundancy in neural activity [5]. This has led to the increasingly popular perspective that collective spiking events in the brain may be described by relatively few underlying factors [2, 6]. Common techniques span simple linear approaches to computationally-intensive deep learning approaches [7, 8]. Importantly, all of these methods, at their core, attempt to extract latent patterns from large and highly complex datasets to help better understand the underlying structure.

Information theory is a subfield of mathematics that aims to quantify, process, and transmit information [9, 10]. Information-theoretic approaches can be appealing because they do not necessarily require or assume the statistical distribution of a dataset, and therefore can be universally applied to all types of data. While the roots of the field are in communications engineering, information-theoretic approaches have been applied across many fields, from quantum physics to molecular dynamics [11, 12]. Within neuroscience, information theory provides a particularly intriguing framework for understanding how information is encoded and communicated in the brain, and has been widely applied to different data types and scientific hypotheses over the past three decades [13–18]. Information-theoretic techniques for neuroscience encompass a large family of methods, all essentially based on determining the information content, or entropy, of neural data [19]. Thus, they are easily generalizable and have been used to describe neural activity at all levels, from single-neuron characteristics to network-wide phenomena [13, 20–22]. A recent information-theoretic application to neural data leverages data compression, specifically the *inverse compression ratio (ICR)*, to evaluate seizure onset in patients with epilepsy, outperforming single-neuron quantitative EEG methods and reducing the reliance on experts for seizure detection [23].

Compression algorithms seek to encode data in a manner that minimizes occupied storage while maximizing, if not totally preserving, the information content of the data. These algorithms are used extensively and ubiquitously in day-to-day digital operations. There are two main types of compression algorithms: lossless, which perfectly encodes a signal, and lossy, which encodes a signal imperfectly in order to balance data quality and size [24, 25]. Compression algorithms can also be divided into temporally-independent methods, which do not use temporal structure to compress the data, and temporally-dependent methods, which do use chronological structure to result in reduced sizes (also known as interframe redundancy) [24]. Common examples of temporally-dependent compressors are video compressors, which often take advantage of between-frame repetitions [26]. Similar to traditional methods used to analyze neural data, compression, especially lossy schemes, can be viewed as a way to distill data into its underlying “essential” components. Furthermore, ICR provides an unbiased upper bound of joint entropy for neural datasets, which can be both temporally and spatially dense [23]. Previous applications of compression to spiking neural data have largely been limited to algorithms designed purely to optimize data storage [27–29]. However, preliminary work has leveraged algorithms with origins in data compression to interpret neural data. Notably, Lempel-Ziv complexity, a metric closely related to compression ratio, has been used to estimate the entropy rate of many different types of neural signals [30–33]. Lempel-Ziv complexity is further used to calculate the perturbational complexity index (PCI), a metric that has been used to evaluate a subject’s consciousness in combined transcranial magnetic stimulation and electroencephalography (TMS-EEG) experiments [34–36]. Other work has used compression and other closely-related algorithms as a tool to quantify and characterize different features across neuroscience, such as neuron synapses, epileptic seizure detection, and patch-clamp recordings [37–39]. However, compression and its range of variations (lossless, lossy; temporally-dependent, temporally-independent) have yet to be directly characterized as a model-free, convenient, and computationally efficient tool to analyze array-based spiking neural data.

This work evaluates the performance of lossless (gzip) [40] and lossy (H.264, MPEG-2) [26, 41, 42] compression against traditional metrics evaluating spiking neural data, such as average firing rates and Fano factor [43, 44]. It also evaluates the performance of compression against common dimensionality reduction schemes, such as principal component analysis (PCA) and factor analysis (FA) [2]. All seven of these methods (gzip ICR, H.264 ICR, MPEG-2 ICR, average firing rates, Fano factor, PCA, FA) are applied to spiking neural data collected from the primary motor cortex (M1) of two rhesus macaques (U and H) performing simple, highly practiced reaching tasks over six days. Additionally, this work investigates how the lossiness (quantified by bitrate) of H.264 and MPEG-2 corresponds to overall performance. Finally, this work quantifies and compares the computational complexity of all methods used. This work aims to establish evidence for the utility of compression-based tools to interpret spiking neural data.

## 3 Methods

### Recordings

All arrays used in this work were multielectrode Utah arrays (Blackrock Neurotech, Salt Lake City, UT). Monkey U (age 11) was implanted with three 96-channel arrays; two in M1 and one in dorsal premotor cortex (PMd). U’s lateral M1 array was used for electrolytic lesioning experiments; the medial array was not used due to poor signal quality. Monkey H (age 14) was implanted with two 96-channel Utah arrays in M1 and PMd. H’s M1 array was also used for electrolytic lesioning experiments. Monkey U’s arrays were manufactured with an iridium oxide metal coating, while H’s arrays were manufactured with platinum coatings. Monkey H’s arrays were implanted for 2242 days (electrolytic lesions on days 2088, 2129, 2136, 2164, 2172, 2180, 2187, 2192, 2221, 2228). Monkey U’s arrays were implanted for 2680 days (electrolytic lesions on days 1215, 1250, 1264, 1286). To avoid potential overlapping effects between lesions, only lesions with at least 140 successful trials and 2 weeks of time between lesions were selected for use in this work (all four lesions for Monkey U; three lesions on days 2129, 2164, 2221 for Monkey H). All animal procedures were reviewed and approved by Stanford University’s Institutional Animal Care and Use Committee.

### Behavioral Task

Monkeys performed a reaching task during daily recording sessions scheduled approximately every 24 hours, seven days a week. Daily data acquisition consisted of a first recording session, followed by a sham lesion, then a second recording session. The sham lesion period was about 10-15 minutes and was intended to prevent monkeys from learning to differentiate between general recording days versus those with a lesion procedure. On a typical day, animals remained highly engaged during session 1 trials, usually completing hundred of successful reaches. Participation in the reaching task was voluntary; if a monkey failed to actively play the task after being presented with several opportunities to participate, he was returned to his home environment. All data used in this analysis was from session 1 of days where the monkey actively participated in the task (see following *Data Selection* section).

To track reach kinematics, a reflective bead was comfortably secured to the third and fourth digits of the hand (proximal phalanges) and optically tracked at 60 Hz in three dimensions with an infrared camera (Polaris Spectra, Northern Digital Inc., Waterloo, Canada). The bead’s position was projected onto a computer screen as a task cursor and aligned in a virtual workspace so that when the animal extended his arm straight out, the cursor mapped to the origin (0,0). This ensured centered visual alignment without having to reach across the body.

Behavioral reaching tasks are well-known and widely-used to evaluate activity in motor cortex [45–47]. For a single trial of the experimental task, illustrated in Figure 1a, the monkey was trained to move its arm to place and hold the cursor over a green target (12 mm diameter) centered at the origin of the virtual workspace. The hold time was 350 ms for Monkey H and 250 ms for Monkey U due to individual performance differences. Next, one of eight radial targets, evenly spaced at a fixed interval of *π*/4 around an invisible circle with a radius of 100 mm, appeared on the workspace. These radial targets appeared randomly within repeated cycles of the eight target set to ensure even trial counts while avoiding potential confounds due to prediction of target location. The monkey then moved their arm to guide the cursor to the radial target, and again held the cursor over the target. Successful navigation of the cursor from center to the target and holding of the cursor to the target earned a juice reward. The target then reset to the center, beginning the next trial of the task. A successful trial is defined as a trial where the monkey was able to successfully complete all aspects of the task (initial center target hold, reach and hold to the correct radial target) within a several second timeout. Both monkeys were trained to be very familiar and comfortable with the task prior to the collection of any data used in this work. Sessions with Monkey H omitted the bottom and bottom-left targets due to a visual field deficit that presented during initial task training.

**Figure 1.**
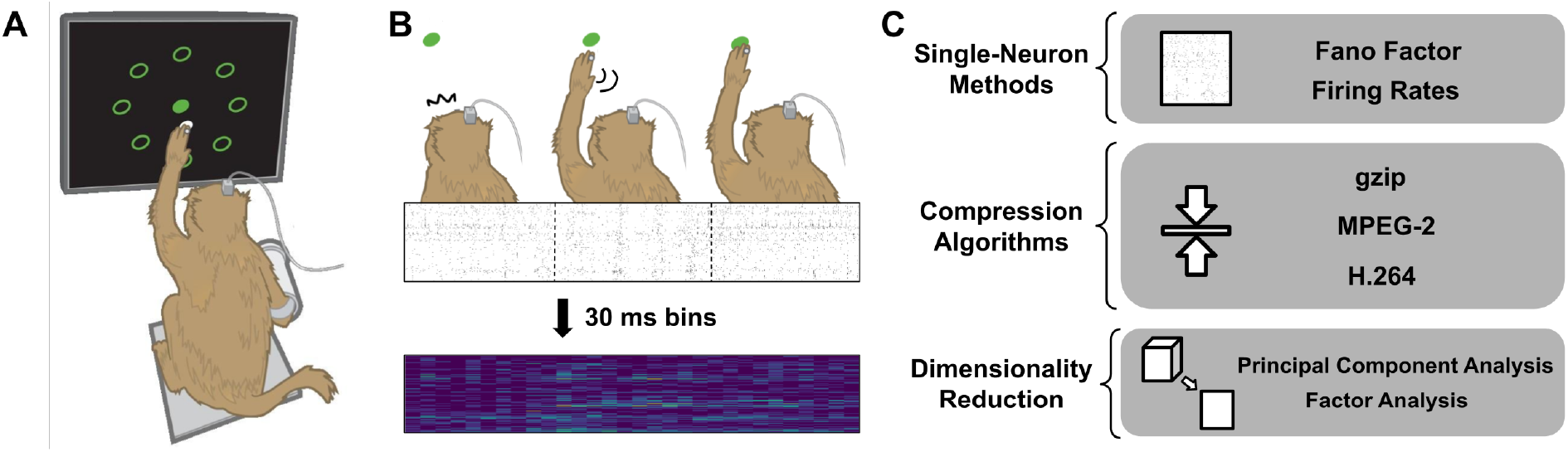
Overview of behavioral task and data analysis. A: Activity was recorded from a 96-channel Utah array implanted in primary motor cortex (M1) of monkeys performing a daily reaching task. Bottom and bottom-left targets were omitted for monkey H due to a visual deficit. All eight radial targets are shown in this figure for illustrative purposes. B: For each trial, 300 ms non-overlapping spike rasters, centered at subject reaction time, maximum arm velocity, and target acquisition time, were concatenated and binned at 30 ms. C: The resulting binned dataset was then analyzed with two single-neuron neuroscience metrics: Fano factor and average firing rate; inverse compression ratio (ICR) under three different compression algorithms: lossless gzip, lossy H.264, and lossy MPEG-2; and two dimensionality reduction techniques: principal component analysis and factor analysis.

### Electrolytic Lesioning

On lesioning days, electrolytic lesions were performed through the recording array during the break period between session 1 and session 2. For both scientific and ethical reasons, lesions were intended to alter an animal’s reach performance but leave overall mobility intact, resulting in a minimal number of failed task trials or frustration. Lesioning procedures and parameters are available in previously published work [48]. The first three days of each dataset was collected with no neural perturbation, and the last three days of the dataset was collected after administering a small electrolytic lesion to M1.

### Data Selection

As previously stated, this work analyzes recordings from three days immediately before the lesioning day and three days immediately after the lesioning day. This data was used for this study due to the clear and large perturbation that electrolytic lesion causes to the the area of cortex we record from. Only successful trials from session 1 of each day are included in this analysis; due to individual differences in successful performance, 500 trials were included for each of Monkey U’s recording sessions across his four lesions, while 200 trials were included for each of Monkey H’s recording sessions across his first and third lesions, and 140 trials were included for each of his recording sessions for his second lesion. From each trial, three non-overlapping epochs of time were extracted: 300 ms centered around the monkey’s reaction to the appearance of the radial target, 300 ms centered around the time at which the monkey’s arm reached maximum velocity, and 300 ms centered around the time at which the monkey successfully acquired the target, as shown in Figure 1b. The resulting recordings were chronologically stitched together to create a 900 ms selection of data per trial and thresholded at -3.5 times the RMS of the signal to extract spiking events. Additionally, all spiking events were binned at 30 ms bins before further analysis.

### Data Analysis

To evaluate the performance of ICR in detecting changes in neural data as a result of electrolytic lesion, seven different metrics were calculated, detailed below. An overview of the analyses is available in Figure 1c. All code was written in Python (version 3.10.6). MPEG-2 encoding was performed using the FFmpeg software package (version 4.4.2-0ubuntu0.22.04.1), and H.264 encoding was performed with the additional software library libx264-163 (version 2:0.163.3060+git5db6aa6-2build1). All code was excuted in the Brain Interfacing lab’s computational environment container (https://github.com/bil/comp-env).

#### Single-Neuron Methods

Two common methods of analyzing neural data, Fano factor and average firing rates, were calculated for each 900 ms selection of data from each trial. These metrics are labeled as “single-neuron” because they can be calculated with single neurons/channels alone and without direct consideration of population-level trends. Furthermore, these two methods were selected not only because of their frequent use in neuroscience research but also because they are often used to approximate neural activity. The Fano factor was calculated as the variance over the mean of the spiking data per trial [44]:

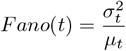

Average firing rates were calculated by summing total spikes over total time:

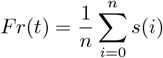

Where *n* = 900 ms, *s*(*i*) is whether or not a neuron spiked at a given time *i* within the selection, and *t* is the trial index.

#### Dimensionality Reduction

Two linear dimensionality reduction analyses, principal component analysis (PCA) and factor analysis (FA), were selected to evaluate the performance of ICR against typical population analyses. PCA and FA are both commonly used in neuroscience research to evaluate population-level trends in neural activity. Similar to data compression, these dimensionality reduction methods aim to exploit statistical characteristics of the data to extract meaningful features. However, unlike data compression, these methods require linearity assumptions and are furthermore not designed for efficient use with very large datasets. Each of the trials were concatenated, forming one matrix (*R*^*c×sn*^, where *c* is the number of channels, *s* is the number of successful trials used, and *n* is 30 the number of bins per trial (30 at 30 ms). These binned, concatenated matrices were each used for these dimensionality reduction analyses. PCA was performed with *n* = 96 desired components, and FA was performed with *n* = 10 desired factors. These analyses were performed using the *PCA* and *FactorAnalysis* modules from the *scikit-learn* library for Python. This work primarily examines the total explained variance in the top factors identified using both of these methods as a way to estimate the structural complexity of the system; more variance in fewer dimensions suggests a more simplistic system than a greater spread of variance over a large number of dimensions. For FA, the common variance, also known as communality, is used as explained variance.

Because only one set of explained variances can be generated per instance of PCA/FA, results from dimensionality reduction may lack statistical power if the entire dataset is used to make comparisons. In order to generate statistically comparable measurements between dimensionality reduction algorithms and other metrics, a bootstrapped analysis was performed as follows. One-fourth of the original successful trial set was selected without replacement. This subset was then used to generate total explained variances with PCA/FA. The top three explained variances were then summed. While there is no standard number of top factors/components to use in estimating population variance from dimensionality reduction, previous work shows that the top three PCs are generally sufficient to explain spikes [49]. This process was repeated until *n* number of explained variance sums were generated, where *n* is the number of trials in the original successful trial set (other metrics are calculated per-trial instead of over multiple trials). This bootstrapped set of explained variance sums was used for all statistical testing.

#### Compression Algorithms

At first glance, many other information-theoretic metrics, such as entropy, mutual information, and joint entropy, might make more sense than compression for analytical purposes as they are directly calculated from data. However, for high-dimensional datasets like neural data, computational costs can scale and become overwhelming when attempting to directly calculate these metrics. For example, for an *n*-dimensional dataset, mutual information must be calculated 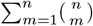 times [23]. This problem is exacerbated as the number of electrodes researchers can directly record from continues to increase, subsequently increasing the dimensionality of recorded data [1].

Entropy is defined as the theoretical minimum number of bits needed to accurately describe a signal. The compressed size (CS) of an object can be interpreted as a practical minimum number of bits needed to accurately describe a signal. Thus, CS serves as an upper bound for the true entropy of a signal. Similarly, the ICR can approximate an upper bound for the entropy of any given dataset in proportion to original raw data size (RS) as follows [23]:

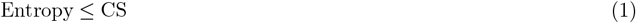

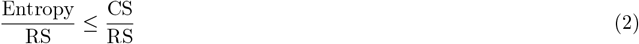

ICR is calculated by comparing compressed and original data sizes:

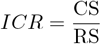

In this work, three different compression algorithms (lossless: gzip, lossy: H.264, MPEG-2) were used to calculate ICR values for each 900 ms selection of data from each trial. All three algorithms were applied to binned spike trains. An explanation of each algorithm is provided below.

gzip is a temporally-independent lossless compression algorithm that is based on the DEFLATE algorithm, which combines the Lempel-Ziv 1 (LZ77) algorithm to first develop a repetition-based code for the data and Huffman coding to secondly compress the coded data even further [40, 50]. LZ77 coding works by sequentially scanning the inputted datastream. Once a section of the datastream is identified as a repetetion of a previous section, it is replaced by a pointer to that previous section [51]. The Huffman coding step takes in this stream of data and pointers from LZ77. It then creates another code that assigns the most repetitive sections of the stream to shorter encodings (and less repetitive sections with longer encodings), so chunks of data/pointers can be described with a number of bytes inversely proportional to their frequency [52]. In other words, the LZ77 step reduces data size based on redundant data, while the Huffman coding step reduces data size based on frequency of redundant data.

H.264 and MPEG-2 are temporally-dependent lossy compression algorithms originally developed for video data. This means that unlike the temporally-independent lossless scheme of gzip, they advantageously look to minimize both spatial and temporal redundancies since they know to expect ordered, 2-dimensional data. This makes these algorithms well-suited to analyzing temporally-ordered recordings formatted in the spatial layout of the physical recording array. All video compressors typically work by exploiting three main types of redundancy: spatial/temporal, based on knowledge about the dependence between consecutive frames and pixels; entropy, based on bit redundancy; and psychovisual, based on knowledge about how the human eye processes visual data [53, 54]. At a high level, MPEG-2 works by first reducing temporal redundancy with interframe prediction, which allows for data savings by reporting only the differences between predicted and real frames [55]. Interframe prediction occurs locally using sets of frames, allowing groups of frames with low activity/change between frames to be more efficiently coded. In MPEG-2, this prediction can occur bidirectionally, where error-prediction frames can be generated from upstream and downstream frames, to optimize output. This generates error-prediction frames that are then separated into 8×8 pixel blocks and projected from the time domain into the frequency domain with the discrete cosine transform (DCT). The DCT maps the data into a series of cosine functions with coefficients, and these coefficients are quantized to further reduce data size. Finally, the quantized DCT coefficients are passed through variable-length coding, where more frequent data blocks are encoded with shorter codenames than rarer data blocks (Huffman coding is a type of variable-length coding). While H.264 is a proprietary and patented algorithm, this work utilizes the open-source libx264 software library, and it is largely based on the same error-prediction and quantized DCT design. However, H.264 takes advantage of additional modern techniques in image and video compression, including (but not limited to) the enabling of multiple reference frames during interframe prediction, flexible options for pixel blocking, and a simplified version of the DCT known as the integer DCT [56, 57]. These changes allow for more efficient encoding on both software and hardware.

For the purposes of these two algorithms, electrode data was arranged according to the grid layout of the Utah array rather than arranged linearly. ICR for H.264 and MPEG-2 were calculated using packet sizes on a per-frame basis, and averaged to determine per-trial average ICR values.

Finally, another feature of temporally-dependent compression algorithms like H.264 and MPEG-2 is bitrate, or number of bits per second that the algorithm can use to describe the data. This is especially interesting in the case of lossy compression, where bitrate can be thought of as a way to coarsely control the amount of data preserved, and therefore also the amount of data lost. Typically, lossy video compressors like H.264 and MPEG-2 use a variable bitrate (VBR) to optimize the visual coherence of the output video. For example, action-packed scenes may result in more complex frames, to which more bits (and thereby a higher VBR) may be dynamically allocated. However, bitrate can also be set to a fixed value, also known as setting a constant bitrate (CBR), in order to allow for more control over the behavior of the algorithm. This work sweeps across CBR ranges, empirically selected for each compression algorithm, to explore the effect of loss on ICR. Each CBR range was selected by determining a consistent low CBR value and high CBR value for each lossy compression algorithm, between which a sharp change in ICR was observed across all lesions and subjects, shown in Figure 3 and Figure 4). This low and high value was then subsampled such that 31 incremental values were evaluated per sweep. CBR ranges differed between compression algorithms, but were invariant between lesions and subjects.

**Figure 2.**
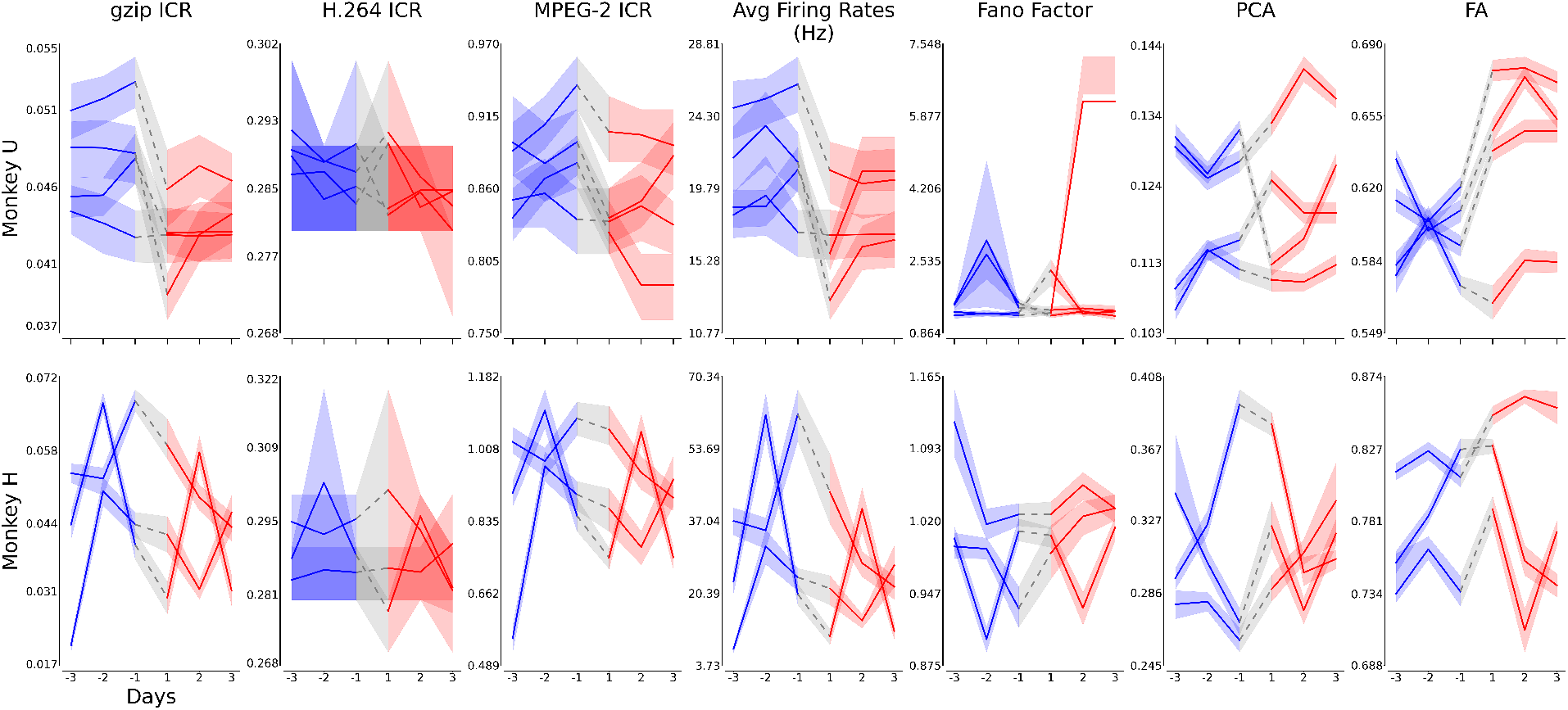
Summary plots for the metrics described in Figure 1 for Monkeys U and H. Each plot chronologically shows the values of the corresponding metric and subject for three pre-lesion days (in blue) and three post-lesion days (in red). The gray region shows the difference between consecutive pre- and post-lesion days. Each line represents a different lesion (U = 4 lesions, H = 3). The first three columns plot the mean and 95^th^ percentile intervals of all gzip inverse compression ratio (ICR), H.264 ICR, and MPEG-2 ICR values. Similarly, the next two columns plot the mean and 95^th^ percentile intervals of all average firing rates and Fano factors. The final two columns plot the total sums of the fractional explained variance for the top 3 factors extracted with principal component analysis (PCA) and factor analysis (FA) on bootstrapped data (see *Methods*). Note that ICR, Fano Factor, and fractional explained variance for PCA and FA are all dimensionless (unitless). These plots highlight the visual quantitative differences between pre- and post-lesion days, as well as intra-day stability, for each metric.

**Figure 3.**
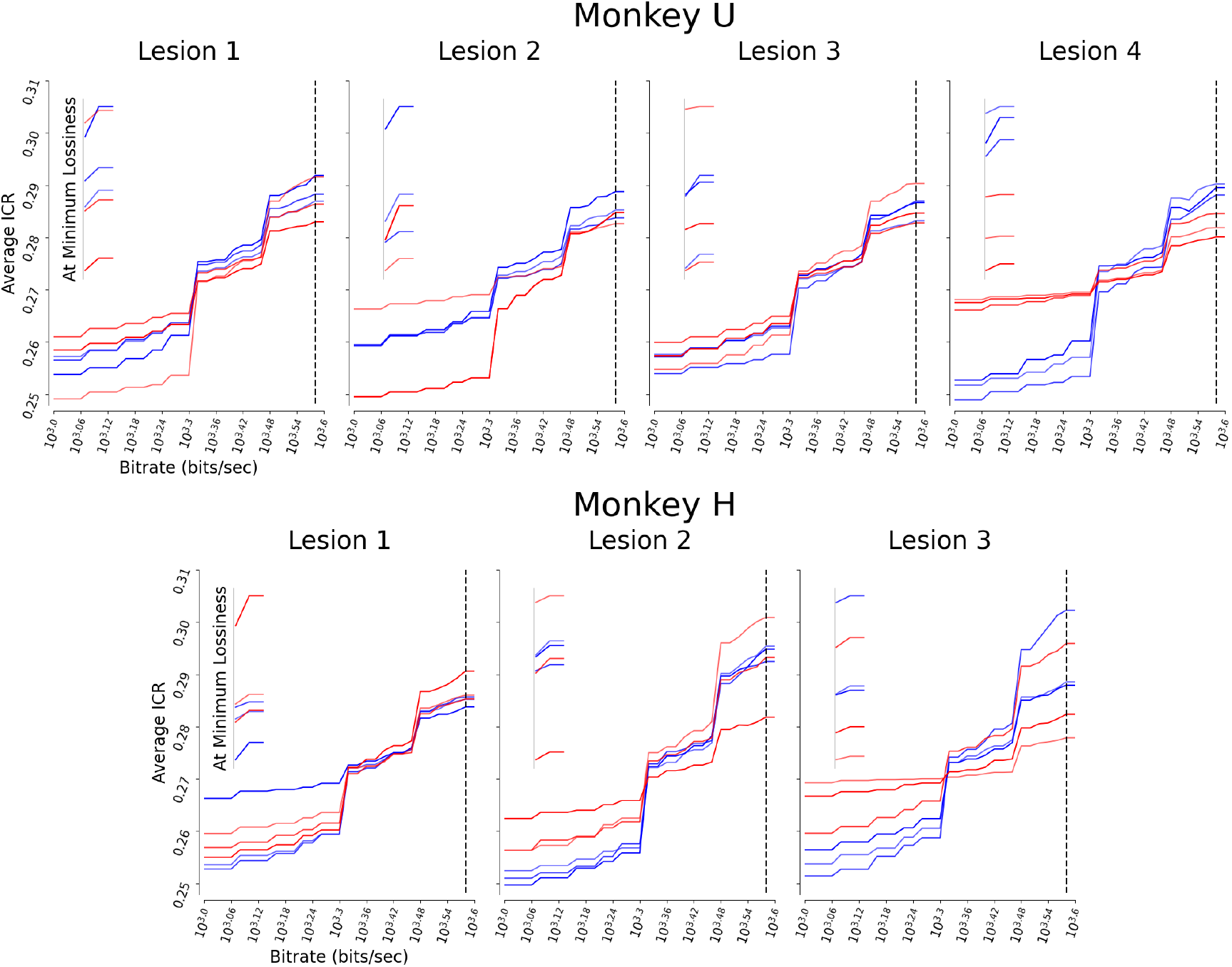
The average ICR after H.264 lossy compression at different constant bitrate (CBR) values. In lossy compression, CBR defines how much data per second the algorithm can use to define outputted data; the higher the bitrate, the lower the amount of data that is lost due to compression. Pre-lesion days are in blue, and post-lesion days are in red. The inset of each graph zooms in on the final three CBR values (marked by the black dotted line), to provide a clearer visual picture of what happens at low data loss. At these final three CBR values, lossiness is minimized due to high bitrates. Note that since this inset is provided for visualization purposes, the axes and scale of each are not the same. Across lesions, we generally see that blue pre-lesion days stratify above red post-lesion days as bitrates get higher and data loss gets lower. The lower bound of this CBR range was empirically selected such that it was the minimum bitrate needed for H.264 to be able to compress the data, and the upper bound was selected such that the reconstruction error (mean squared error) between the original and compressed data no longer improved significantly with increasing bitrate. The specific CBR range selected for H.264 was {10^3^, 10^3.02^, 10^3.04^, 10^3.06^, 10^3.08^, 10^3.1^, 10^3.12^, 10^3.14^, 10^3.16^, 10^3.18^, 10^3.2^, 10^3.22^, 10^3.24^, 10^3.26^, 10^3.28^, 10^3.3^, 10^3.32^, 10^3.34^, 10^3.36^, 10^3.38^, 10^3.4^, 10^3.42^, 10^3.44^, 10^3.46^, 10^3.48^, 10^3.5^, 10^3.52^, 10^3.54^, 10^3.56^, 10^3.58^, 10^3.6^ *}* bits/sec. Note that all x and y-axes are at the same scale and carry over between main plots. Finally, note that ICR (and thus average ICR) is a dimensionless metric.

**Figure 4.**
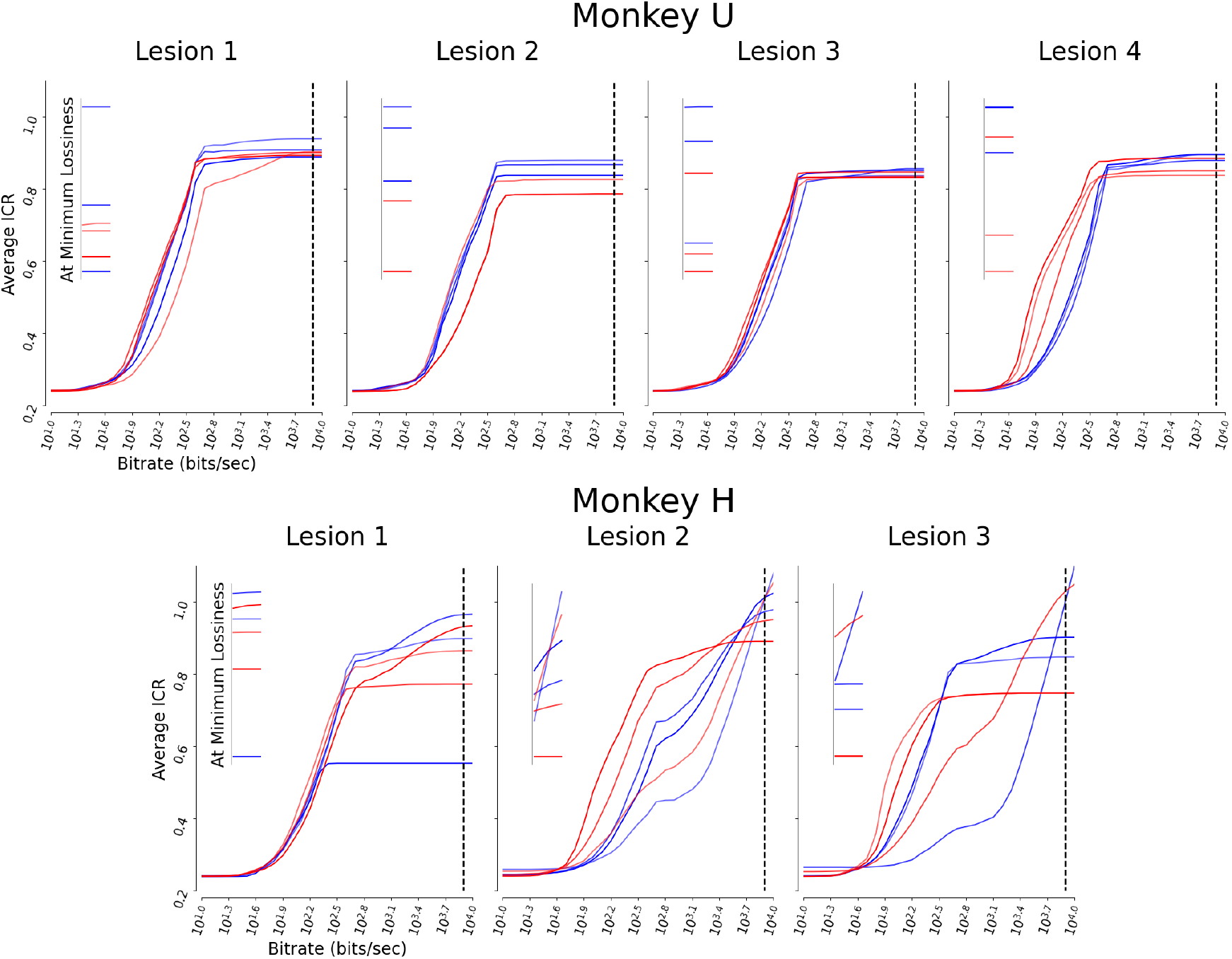
The average ICR after MPEG-2 lossy compression at different constant bitrate (CBR) values. Again, in lossy compression, CBR approximately defines how much data is lost due to compression. Formatted exactly like Figure 3, pre-lesion days are in blue, and post-lesion days are in red. The inset of each graph zooms in on the final three CBR values (marked by the black dotted line), to provide a clearer visual picture of what happens at low data loss. Note that since this inset is provided for visualization purposes, the axes and scale of each are not the same. Similar to H.264 results, we generally see that blue pre-lesion days stratify above red post-lesion days as bitrates get higher and data loss gets lower. The lower bound of this CBR range was selected such that it was the minimum bitrate needed for MPEG-2 to be able to compress the data, and the upper bound was selected such that the reconstruction error (mean squared error) between the original and compressed data no longer improved significantly with increasing bitrate. The specific CBR range selected for MPEG-2 was {10^1^, 10^1.1^, 10^1.2^, 10^1.3^, 10^1.4^, 10^1.5^, 10^1.6^, 10^1.7^, 10^1.8^, 10^1.9^, 10^2^, 10^2.1^, 10^2.2^, 10^2.3^, 10^2.4^, 10^2.5^, 10^2.6^, 10^2.7^, 10^2.8^, 10^2.9^, 10^3^, 10^3.1^, 10^3.2^, 10^3.3^, 10^3.4^, 10^3.5^, 10^3.6^, 10^3.7^, 10^3.8^, 10^3.9^, 10^4^, 10^4.1^ } bits/sec. Note that all x and y-axes are at the same scale and carry over between plots. Note also that average ICR may be > 1 in cases where compressor overhead make the outputted file larger than the input file; this may occur at very high bitrates when compression algorithms are not being used as intended. Finally, note that ICR (and thus average ICR) is a dimensionless metric.

**Figure 5.**
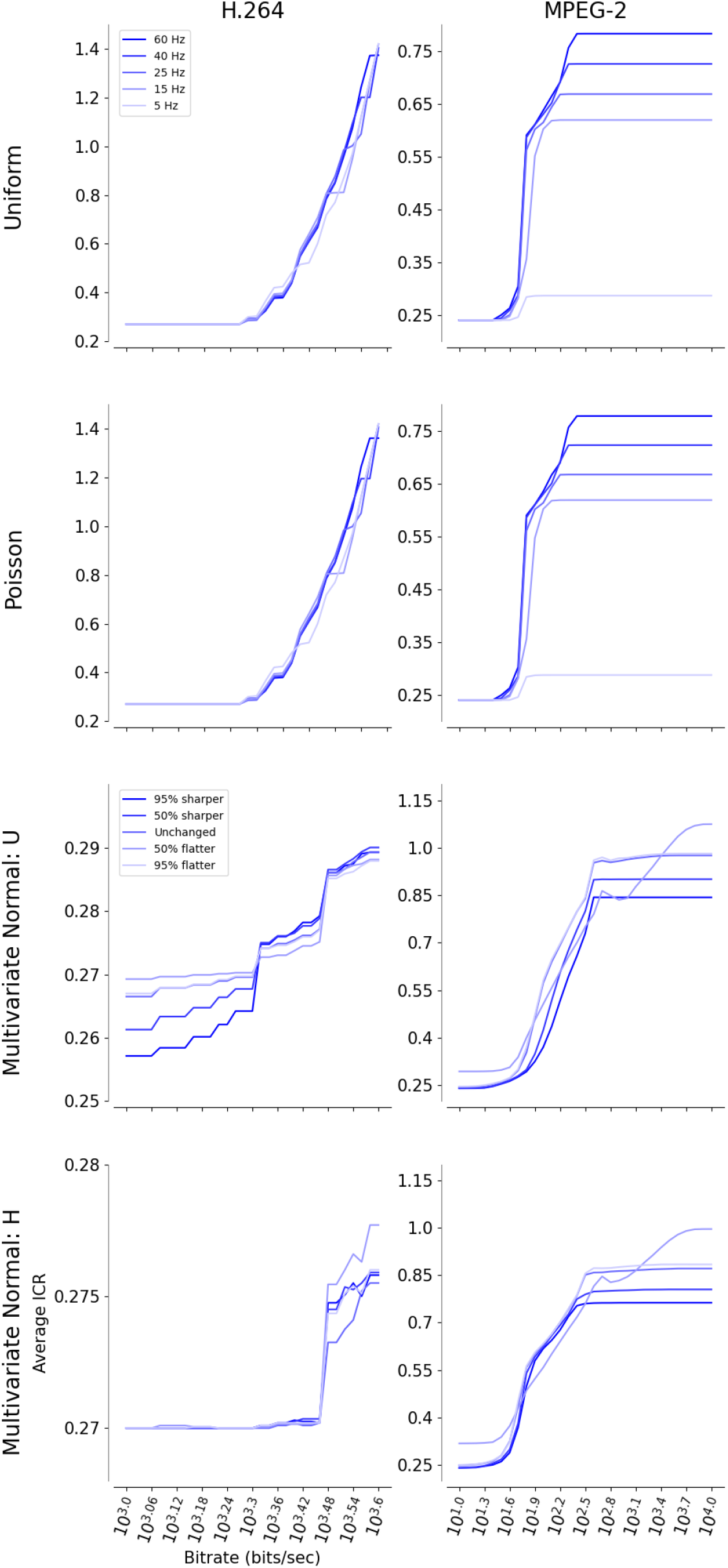
(Previous page.) Simulation results. The leftmost column displays all CBR sweep results for H.264, and rightmost column displays CBR sweep results for MPEG-2. The same CBR ranges used for real neural data were used to generate the displayed plots. The top row shows average ICR values from uniformly-distributed spiking neural data. The second row shows average ICR values from Poisson-distributed spiking neural data. The legend in the upper-left of the first row, defining the different firing frequencies, applies to these first four plots. The third and fourth rows show average ICR values from data drawn from multivariate normal distributions based on a random experimental session from subject U and H, respectively. The legend in the upper-left of the third row, defining the change to the eigenspectra underlying the multivariate normal distribution, applies to these last four plots. Specific details on simulated data are available in paper Methods. Note that all x-axes are at the same scale and carry over between plots. All y-axes plot average ICR, but have different ranges (as labeled). Note also that average ICR may be *>* 1 in cases where compressor overhead make the outputted file larger than the input file; this may occur at very high bitrates when compression algorithms are not being used as intended. Finally, note that ICR (and thus average ICR) is a dimensionless metric.

### Control Simulations

To better understand the relationship between CBR and underlying structure in compressed data, CBR sweeps were also conducted for three sets of simulated datasets. The first two datasets explore how changes in channel spiking rates affect ICR, while the third aims to explore overall shifts in the correlational structure of data. The first set consisted of uniformly sampled spike counts at a given average firing rate (5, 15, 25, 40, 60 Hz). The second set consisted of Poisson-distributed spike counts at a given average firing rate (5, 15, 25, 40, 60 Hz). Finally, the third set consisted of binned spike counts generated by computing and adjusting the eigenspectra of experimental data. All simulated data were presented as 30 ms bins, exactly as experimental data was processed before analysis. Specifically, one experimental session was selected from each subject. The data from each experimental session was then used to calculate a mean and a covariance matrix. The eigenvalues of these covariance matrices were sorted, and the eigenspectra was subsequently flattened or sharpened with respect to a given factor. Explicitly, two exponential functions of form *y* = *Ae*^*cBx*^ were fitted to the eigenspectra derived from neural data from a randomly-selected day from each of the two subjects, where *A* and *B* are parameters derived from fitting. The final parameter, *c* = {− 0.95, − 0.5, 0, 0.5, 0.95}, was chosen to scale the flatness and sharpness of the resulting function. These adjusted eigenspectra was then used to re-calculate covariance matrices, which were in turn used with the mean to define the two (per-subject) multivariate normal distributions. These distributions were then sampled and clipped to a range of 0-6, in order to stay biologically plausible (200 Hz maximum frequency * 30 ms bin size /1000 ms per second = 6 spikes per bin). Respectively, these four simulations represent a very simple structure-free (and thereby entropically maximal) case, a very statistically structured case, and two very statistically structured and biologically inspired cases.

## 4 Results

Generally, as shown in Figure 2, all three ICR methods calculated higher values for the pre-lesion days than the post-lesion days, suggesting that the pre-lesion recordings are more statistically complex than the post-lesion recordings. This is similarly reflected in the plot of average firing rates, as shown in Figure 2. Fano factor generally seems to weakly invert the trend, showing higher metrics calculated after lesioning. Because Fano factor is a measure of variability, this may indicate that post-lesion data tends to have a wider dispersion in values. When taken alongside our ICR results, which find that post-lesion data is more compressible, this may in turn mean that post-lesion data tends to be noisier; in other words, post-lesion data may carry more shared noise artifacts that do not carry much neurally-relevant information but still affect the recorded signals. Finally, we see that the total explained variance for the top three principal components/factors identified by PCA and FA, respectively, generally increases after lesioning. This again may indicate that pre-lesion data is more structurally complex than post-lesion data, as the system condenses its activity to a smaller number of dimensions.

These trends were then statistically quantified using the two-sided Mann-Whitney U test comparing calculated metrics from all three pre-lesion days against all three post-lesion days. ICR methods significantly (*p <* 0.01) detected lesions with 85.7% accuracy (gzip: 85.7%, H.264: 85.7% MPEG-2: 85.7 %), single-neuron methods detected lesions with 78.6% accuracy (average firing rates: 71.4%, Fano factor: 85.7%), and the variance in the top three features detected by dimensionality reduction techniques detected lesions with 100% accuracy. Here, we define accuracy as the number of times the metric changes statistically significantly when a lesion is administered. Note that while many of these accuracy percentages are coincidentally similar, they do not fail to detect the same consistent lesions (see Supplementary Table 1). Additionally, note that these dimensionality reduction tests were done on bootstrapped metrics.

The Levene test (using mean as the center measure) was also used to evaluate changes in measure stability before and after lesion using this same pooled dataset. Significantly (*p <* 0.01) dissimilar variances were identified for 66.7% of lesions with ICR methods (gzip: 71.4%, H.264: 57.1%, MPEG-2: 71.4%), 92.9% of lesions with single-neuron methods (average firing rates: 100%, Fano factor: 85.7%), and 85.7% of lesions with the summed top three features detected by dimensionality reduction techniques (PCA: 85.7%, FA: 85.7%). Again, note that these dimensionality reduction tests were done on bootstrapped metrics. The results of all statistical tests are available in Supplemental Table 1.

The CBR was additionally fixed and swept for both lossy compression algorithms to investigate the relationship between ICR and compressor bitrate. The results for H.264 are plotted in Figure 3, and the results for MPEG-2 are plotted in Figure 4. As shown in plot insets, pre-lesion days (blue) tend to have greater average ICR values than post-lesion days (in red) when the compressors are able to operate with high bitrates (minimizing loss), though there is considerable heterogeneity. Interestingly, this trend does not seem to emerge or remain stable until compression algorithms reach higher bitrates. In fact, for certain lesions, pre-lesion days have reliably lower ICR values than post-lesion days until high bitrates are achieved. These trends are true for both H.264 and MPEG-2 ICR values. Additionally, some MPEG-2 ICR values calculated from this data are occasionally greater than 1. This can be attributed to the addition of compressor-specific formatting overhead, and is fairly common in compressing small video or image files that may already have low redundancy.

Results of control simulations are plotted in Figure 8, which shows the average ICR against the same CBR ranges as previous sweeps for each simulated dataset. While results are extremely similar between uniformly-distributed and Poisson-distributed spiking data, the exact values are not identical.

Additionally, this work quantified the computational complexity of all seven metrics by recording the average elapsed time it took for each metric to be calculated. Ten random experiment sessions were chosen without replacement for this comparison. All seven metrics were computed using identically-structured data. The results are shown below in Table 1.

**Table 1:** Average elapsed time in seconds for each of the seven metrics evaluated in this work, using ten random experiment sessions used to generate key results in this paper.

| Metric | Average Elapsed Time (sec) |
| --- | --- |
| Average Firing Rates | 0.00115 |
| Fano Factor | 0.00301 |
| gzip ICR | 1.20 |
| H.264 ICR | 38.9 |
| MPEG-2 ICR | 38.2 |
| PCA | 0.0278 |
| FA | 0.280 |

Another important consideration when quantifying computational complexity is how each algorithm scales as the dataset increases in size. Because compression algorithms’ efficiency is a direct factor of the amount of redundancy in a dataset, larger datasets with inherently more opportunities for redundancy are expected to be far more efficient than other metrics as data gets bigger. To evaluate this, three datasets were generated from 1, 25, and 100 experimental sessions, randomly selected without replacement. Each dataset stacked the channel recordings in 10 x 10, 50 x 50, and 100 x 100 formats reflecting the real layout of the physical arrays. (See also *Compression Algorithms* in Methods section.) Each dataset was evaluated using the same methods as in Table 1. This analysis was also repeated for three similarly-formatted simulated datasets, where spikes were generated under a Poisson distribution with a 25 Hz average firing rate. As the data increases in size, all metrics similarly scale in elapsed time, except for the lossy ICR algorithms, which stay relatively stable. The results are show below in Table 2.

**Table 2:** Elapsed time in seconds for each of the seven metrics evaluated in this work. Real data refers to using 1, 25, and 100 randomly-selected experiment sessions (not necessarily those directly evaluated in the key results of this paper). Simulated data refers to using spike trains generated under a Poisson distribution with an average firing rate of 25 Hz. Spikes were formatted similarly to the 96-channel array recordings, and arranged to reflect the same format as the 1, 25, and 100-session data.

| Metric | Elapsed Time (sec) |  | Data Size (Mb) | Total # Sessions |
| --- | --- | --- | --- | --- |
|  | Real Data | Simulated Data |  |  |
| Avg Firing Rates | 0.00223 | 0.00128 | 11.52 | 1 |
| Avg Firing Rates | 0.0532 | 0.0260 | 288 | 25 |
| Avg Firing Rates | 0.129 | 0.105 | 1152 | 100 |
| Fano Factor | 0.00303 | 0.00451 | 11.52 | 1 |
| Fano Factor | 0.0900 | 0.120 | 288 | 25 |
| Fano Factor | 0.354 | 0.472 | 1152 | 100 |
| gzip ICR | 1.59 | 1.51 | 11.52 | 1 |
| gzip ICR | 44.8 | 44.0 | 288 | 25 |
| gzip ICR | 181 | 177 | 1152 | 100 |
| H.264 ICR | 65.2 | 60.3 | 11.52 | 1 |
| H.264 ICR | 65.8 | 69.2 | 288 | 25 |
| H.264 ICR | 71.7 | 75.6 | 1152 | 100 |
| MPEG-2 ICR | 68.2 | 58.2 | 11.52 | 1 |
| MPEG-2 ICR | 60.8 | 56.4 | 288 | 25 |
| MPEG-2 ICR | 54.5 | 53.8 | 1152 | 100 |
| PCA | 0.0351 | 0.0452 | 11.52 | 1 |
| PCA | 4.83 | 5.27 | 288 | 25 |
| PCA | 218 | 223 | 1152 | 100 |
| FA | 0.431 | 0.165 | 11.52 | 1 |
| FA | 4.19 | 2.28 | 288 | 25 |
| FA | 6.94 | 9.29 | 1152 | 100 |

## 5 Discussion

This paper is an empirical analysis of ICR as a tool for detecting perturbations, like electrolytic lesion, in neural data. In general, all three ICR methods calculate higher values for the pre-lesion days than the acute post-lesion days, suggesting that the pre-lesion recordings are more statistically complex than the post-lesion recordings.

This may provide further evidence for the function of redundant population dynamics in the brain, a hypothesized feature of neural dynamics that has been shown to help the brain maintain stable function even as the system changes over time [58–62]. Previous work has shown that changes to specific neurons, even when highly-tuned and predictive, do not necessarily change overall population dynamics due to redundant computations in the network [60]. Indeed, the animal was able to return to playing the task immediately after the administration of each lesion. While the system might lose specific neurons of varying importance during lesioning, it may be able to safely fall back on redundant computations in its repertoire to support the same physical behaviors [61, 63]. This might indicate that lesioning causes the system to leverage redundancy while decreasing the structural complexity, and therefore the ICR, of the data.

Furthermore, peri-infarct cortex is known to have increased inhibition during recovery after stroke [62, 64, 65]. Lower general activity levels in the lesioned area, highlighted by the drop in average firing rates after lesion, would also explain lower post-lesion ICR values. However, changes in single-neuron metrics like firing rate or even changes in the overall distribution of data alone are not sufficient to explain the large fluctuations we see in ICR before and after lesion. This is evidenced by the control simulations, which show how changes in average firing rate, under both uniform and Poisson distributions, affect ICR. Average firing rates under both distributions do not result in very large differences for H.264, though the compression algorithm is generally able to compress lower firing rates more efficiently (lower ICR) than higher firing rates. The average ICR under MPEG-2 is highly dependent on average firing rates, very clearly compressing lower firing rates more efficiently than higher firing rates. Figure 2 shows how MPEG-2 ICR and average firing rate measures have very similar shapes. On the other hand, large eigenspectra-based structural changes to the data also do not result in very big differences for either H.264 or MPEG-2, though both compressors are able to compress data with sharpened eigenspectra more efficiently at lower bitrates. Thus, ICR may be a reflection some other signature of the perturbation, likely a combination of individual neuron spiking activity, correlational structure, and other types of changes that may result in changing levels of data redundancy. It is notable that MPEG-2, an older and simpler compression algorithm, is much more sensitive to base changes in average compression rate. However, because this work is an empirical study of ICR, the exact underlying mechanisms and features of neural activity that drive these changes in compressibility are not fully understood yet and will be expanded on in future work.

Statistical comparisons of pre- and post-lesion metrics show that ICR more accurately predicts the presence of a lesion than single-neuron methods. However, single-neuron methods were only slightly worse. Additionally, dimensionality reduction methods outperform both ICR and single-neuron methods. Because ICR in this work is only calculated using general-purpose compression algorithms, these results indicate that ICR has potential in accurately detecting neural lesions, and would likely perform even better with algorithms customized to neural data. These statistical comparisons between pre- and post-lesion metrics also show that ICR experiences fewer statistical changes in its population variance than both single-neuron metrics and dimensionality reduction. These results may indicate that ICR values are more stable to system perturbations than these other computational approaches. This reflects the possibility that ICR is able to reflect multiple features that may present themselves as quantifiable information; a combination of changes occurring at different levels in the brain. This may also reflect how ICR is able to capture nonlinear changes more accurately than dimensionality reduction; PCA and FA both rely on linearity assumptions.

The sweeps show that generally, pre-lesion ICR values are higher than post-lesion ICR values when the compressor is allowed to maximize its performance with high bitrates. At low bitrates, lossy compressors often calculate lower ICR values for pre-lesion days than post-lesion days. However, this trend inverts as compressors are allowed higher bitrates. In other words, at low bitrates (high reconstruction error), compressors detect higher entropic information content in neural data than at high bitrates (low reconstruction error). This may indicate the presence of relatively higher uncorrelated noise in post-lesion data. As the system is given more bits to describe the data, it may be better able to represent non-noisy features that reflect the underlying state of the neural system. Notably, modern video compressors like H.264 and MPEG-2 usually leverage variable bitrates instead of constant bitrates, as VBR is able to better handle the changing motion and activity levels inherent to video [42, 66]. This motivates the development of compression algorithms specifically designed for the unique structure and activity patterns of neural data.

Finally, the computational complexity comparison shows that ICR generally takes at least one order of magnitude (if not several) longer to compute than other methods when simply considering small, single-session data. This may partially be attributed to the way each metric is computed; single-neuron and gzip ICR metrics were computed once per trial, H.264 and MPEG-2 ICR metrics were computed per frame (time point, or 30 ms bin) then used to take a median per trial, and FA and PCA were each computed once per experimental session (all trials). The compression algorithms used in this work are not designed for use on analyzing neural data, so it is not surprising that they behave inefficiently. MPEG-2 and H.264 are both video compressors designed to operate on frames with thousands and millions of bytes rather than tens and hundreds. Importantly, the results of the computational complexity evaluation demonstrate that as ICR scales with larger datasets, lossy compression algorithms outcompete other population-based metrics like lossless compression, PCA, and FA, in terms of computational efficiency. As researchers move to higher-electrode channel systems, computationally-efficient unsupervised methods for population analyses of neural data, like lossy ICR, will become more and more necessary for tractable analyses.

Importantly, this empirical work sets the stage for future work that leverages compression algorithms for neuroscientific analysis. There are many additional features of neural data itself that future work can explore. Spike sorting, which theoretically removes random noise from data and makes data more biological by extracting neuron-level activity (instead of activity at the resolution of the recording technology), doesn’t necessarily result in consistent improvement with traditional classifiers [67– 71]. Future work may evaluate whether ICR is similarly robust to spike sorting. Additionally, future work may investigate whether ICR is robust to changes in the neural perturbation and recording modalities. While the data used in this study is limited to electrolytic lesion and array recordings, ICR is easily applied to other experimental settings because it operates on the very bits themselves. In fact, imaging data may very well be better-suited to use with video compression algorithms than electrophysiological data.

This work also motivates the development of new data compression algorithms for complexity estimation and analysis. More broadly, recent work has demonstrated utilizing the Lempel-Ziv compression algorithm to generate sequential probability assignments, a universally-consistent result that demonstrates how novel compression methods may be used to extract features from all types of data [72]. For neuroscience-specific applications, inherent features of neural data, like knowledge of firing rate and region of the brain, can be used to inform both efficiency and robustness of compression algorithms designed for ICR.

Additional work that seeks to characterize and/or improve the underlying noise profile of neural data may further aid the development of these algorithms [73, 74].

## 6 Conclusion

This work sought to introduce data compression as a useful tool for neuroscientific analysis via the inverse compression ratio. Through a survey of two single-neuron methods, three compression algorithms, and two dimensionality reduction techniques applied to NHP data before and after lesion, this work showed that ICR is able to better detect the presence of neural lesions than single-neuron methods, and approaches the accuracy of dimensionality reduction techniques. Furthermore, ICR remains more stable than single-neuron metrics. Additionally, this work investigated the importance of lossy compressor bitrate on ICR, finding that pre- and post-lesion ICR relationships are not preserved across all bitrates, emerging only at higher bitrates with lower reconstruction error. Finally, this work characterized the relative inefficiency of the compression algorithms, prompting the development of compressors specifically designed for neural data analysis.

Thus, data compression may serve as a nascent yet already-ubiquitous computational method to investigate the complexity of neural data, alongside traditional methods like dimensionality reduction. These results motivate future work in developing custom compression algorithms to optimize both the storage and interpretability of neural data as researchers’ window into the brain continues to scale.

## 7 Author Contributions

A Tor was responsible for conceptualization, data curation, formal analysis, investigation, methodology, software, validation, visualization, writing, and review/editing. Y Wu was responsible for conceptualization, data curation, formal analysis, methodology, software, and review/editing. SE Clarke was responsible for data curation, supervision, writing, and review/editing. L Yamada was responsible for methodology, supervision, and review/editing. T Weissman was responsible for conceptualization, methodology, supervision, and review/editing. P Nuyujukian was responsible for conceptualization, funding acquisition, methodology, project administration, resources, supervision, writing, and review/editing. The members of the Brain Interfacing Laboratory provided additional support in review/editing.

## 8 Acknowledgments

We thank S Baker for veterinary support, and K Chin and M Truong for administrative support. The members of the Brain Interfacing Laboratory are Michelle S Wechsler, Mackenzie Risch, Alexandra Paraskevopoulou, Stephen I Ryu, Alissa S Ling, Iliana E Bray, Elizabeth Jun, Michael P Silvernagel, Kenji Y Marshall, Muhammad Abdulla, and Sydney Hunt. MS Wechsler, A Paraskevopoulou, K Lebedev, and MJ Risch were responsible for animal care and surgical support. SI Ryu was responsible for nonhuman primate array implantation. AS Ling, IE Bray, E Jun, MP Silvernagel, K Marshall, MU Abdulla, and S Hunt assisted in animal care. A Tor was supported by the Department of Defense (DoD) through the National Defense Science & Engineering Graduate (NDSEG) Fellowship Program and by a training grant from the National Science Foundation (1828993). SE Clarke was supported by a Stanford School of Medicine’s Dean’s Posdoctoral Fellowship. This work was supported by a Stanford Human-Centered AI Seed Research Grant to SE Clarke and P Nuyujukian. This work was additionally supported by the National Institutes of Health (R01NS123517, R01NS130789, U19NS118284) and the Stanford Wu Tsai Neurosciences Institute to P Nuyujukian.

## Figures

Note: All figures updated

**Supplemental Table 1:** Mann-Whitney U test and Levene test statistics and p-values when comparing calculated metrics between pre-lesion and post-lesion days.

| Subject | Lesion | Metric | Test | Statistic | P-Value |
| --- | --- | --- | --- | --- | --- |
| U | 1 | gzip ICR | Mann-Whitney U | 2.061E06 | 0.000E00 |
| U | 1 | gzip ICR | Levene | 5.754E01 | 4.387E-14 |
| U | 1 | H.264 ICR | Mann-Whitney U | 1.233E06 | 1.231E-06 |
| U | 1 | H.264 ICR | Levene | 5.102E-01 | 4.751E-01 |
| U | 1 | MPEG-2 ICR | Mann-Whitney U | 1.355E06 | 2.540E-22 |
| U | 1 | MPEG-2 ICR | Levene | 3.220E00 | 7.283E-02 |
| U | 1 | Firing Rates | Mann-Whitney U | 2.065E06 | 0.000E00 |
| U | 1 | Firing Rates | Levene | 4.685E01 | 9.259E-12 |
| U | 1 | Fano Factor | Mann-Whitney U | 7.172E05 | 3.038E-66 |
| U | 1 | Fano Factor | Levene | 1.768E03 | 3.764E-304 |
| U | 1 | PCA | Mann-Whitney U | 5.773E04 | 0.000E00 |
| U | 1 | PCA | Levene | 1.350E02 | 1.456E-30 |
| U | 1 | FA | Mann-Whitney U | 2.863E04 | 0.000E00 |
| U | 1 | FA | Levene | 9.876E01 | 6.427E-23 |
| U | 2 | gzip ICR | Mann-Whitney U | 1.849E06 | 1.278E-204 |
| U | 2 | gzip ICR | Levene | 3.674E00 | 5.537E-02 |
| U | 2 | H.264 ICR | Mann-Whitney U | 1.221E06 | 7.966E-06 |
| U | 2 | H.264 ICR | Levene | 1.023E01 | 1.399E-03 |
| U | 2 | MPEG-2 ICR | Mann-Whitney U | 1.967E06 | 5.710E-277 |
| U | 2 | MPEG-2 ICR | Levene | 6.511E01 | 1.012E-15 |
| U | 2 | Firing Rates | Mann-Whitney U | 1.142E06 | 4.669E-01 |
| U | 2 | Firing Rates | Levene | 7.260E02 | 2.241E-143 |
| U | 2 | Fano Factor | Mann-Whitney U | 3.863E05 | 6.295E-213 |
| U | 2 | Fano Factor | Levene | 2.765E04 | 0.000E00 |
| U | 2 | PCA | Mann-Whitney U | 5.623E04 | 0.000E00 |
| U | 2 | PCA | Levene | 4.260E02 | 1.322E-88 |
| U | 2 | FA | Mann-Whitney U | 5.000E01 | 0.000E00 |
| U | 2 | FA | Levene | 6.332E01 | 2.471E-15 |
| U | 3 | gzip ICR | Mann-Whitney U | 1.328E06 | 1.251E-17 |
| U | 3 | gzip ICR | Levene | 1.693E01 | 3.976E-05 |
| U | 3 | H.264 ICR | Mann-Whitney U | 1.100E06 | 2.505E-01 |
| U | 3 | H.264 ICR | Levene | 1.046E-01 | 7.464E-01 |
| U | 3 | MPEG-2 ICR | Mann-Whitney U | 1.292E06 | 1.656E-12 |
| U | 3 | MPEG-2 ICR | Levene | 1.068E01 | 1.095E-03 |
| U | 3 | Firing Rates | Mann-Whitney U | 1.441E06 | 1.632E-40 |
| U | 3 | Firing Rates | Levene | 1.215E02 | 9.959E-28 |
| U | 3 | Fano Factor | Mann-Whitney U | 1.577E06 | 7.769E-81 |
| U | 3 | Fano Factor | Levene | 9.853E02 | 2.983E-187 |
| U | 3 | PCA | Mann-Whitney U | 1.325E06 | 3.935E-17 |
| U | 3 | PCA | Levene | 1.499E02 | 1.169E-33 |
| U | 3 | FA | Mann-Whitney U | 1.329E06 | 9.216E-18 |
| U | 3 | FA | Levene | 1.942E02 | 8.150E-43 |
| U | 4 | gzip ICR | Mann-Whitney U | 2.105E06 | 0.000E00 |
| U | 4 | gzip ICR | Levene | 1.302E02 | 1.473E-29 |
| U | 4 | H.264 ICR | Mann-Whitney U | 1.439E06 | 6.474E-45 |
| U | 4 | H.264 ICR | Levene | 1.670E02 | 3.198E-37 |
| U | 4 | MPEG-2 ICR | Mann-Whitney U | 1.591E06 | 2.743E-86 |
| U | 4 | MPEG-2 ICR | Levene | 3.708E00 | 5.426E-02 |
| U | 4 | Firing Rates | Mann-Whitney U | 2.181E06 | 0.000E00 |
| U | 4 | Firing Rates | Levene | 3.825E01 | 7.069E-10 |
| U | 4 | Fano Factor | Mann-Whitney U | 2.130E06 | 0.000E00 |
| U | 4 | Fano Factor | Levene | 1.156E03 | 1.445E-214 |
| U | 4 | PCA | Mann-Whitney U | 2.047E06 | 0.000E00 |
| U | 4 | PCA | Levene | 7.393E02 | 1.070E-145 |
| U | 4 | FA | Mann-Whitney U | 0.000E00 | 0.000E00 |
| U | 4 | FA | Levene | 1.012E02 | 1.935E-23 |
| H | 1 | gzip ICR | Mann-Whitney U | 1.847E05 | 4.317E-01 |
| H | 1 | gzip ICR | Levene | 3.707E02 | 3.520E-72 |
| H | 1 | H.264 ICR | Mann-Whitney U | 1.663E05 | 1.552E-02 |
| H | 1 | H.264 ICR | Levene | 7.657E00 | 5.742E-03 |
| H | 1 | MPEG-2 ICR | Mann-Whitney U | 1.785E05 | 8.023E-01 |
| H | 1 | MPEG-2 ICR | Levene | 4.867E02 | 8.620E-91 |
| H | 1 | Firing Rates | Mann-Whitney U | 1.845E05 | 4.565E-01 |
| H | 1 | Firing Rates | Levene | 1.226E02 | 3.438E-27 |
| H | 1 | Fano Factor | Mann-Whitney U | 1.697E05 | 8.509E-02 |
| H | 1 | Fano Factor | Levene | 9.977E01 | 1.276E-22 |
| H | 1 | PCA | Mann-Whitney U | 1.618E05 | 2.384E-03 |
| H | 1 | PCA | Levene | 1.226E02 | 3.383E-27 |
| H | 1 | FA | Mann-Whitney U | 1.247E05 | 3.419E-20 |
| H | 1 | FA | Levene | 4.125E01 | 1.927E-10 |
| H | 2 | gzip ICR | Mann-Whitney U | 1.331E05 | 2.770E-37 |
| H | 2 | gzip ICR | Levene | 3.667E00 | 5.583E-02 |
| H | 2 | H.264 ICR | Mann-Whitney U | 9.711E04 | 9.139E-03 |
| H | 2 | H.264 ICR | Levene | 1.487E-01 | 6.999E-01 |
| H | 2 | MPEG-2 ICR | Mann-Whitney U | 1.253E05 | 4.259E-26 |
| H | 2 | MPEG-2 ICR | Levene | 3.893E01 | 6.962E-10 |
| H | 2 | Firing Rates | Mann-Whitney U | 1.406E05 | 3.149E-50 |
| H | 2 | Firing Rates | Levene | 1.316E01 | 3.029E-04 |
| H | 2 | Fano Factor | Mann-Whitney U | 5.021E04 | 3.245E-27 |
| H | 2 | Fano Factor | Levene | 1.814E01 | 2.282E-05 |
| H | 2 | PCA | Mann-Whitney U | 1.028E05 | 3.180E-05 |
| H | 2 | PCA | Levene | 1.325E00 | 2.500E-01 |
| H | 2 | FA | Mann-Whitney U | 1.114E05 | 4.147E-11 |
| H | 2 | FA | Levene | 7.190E01 | 1.016E-16 |
| H | 3 | gzip ICR | Mann-Whitney U | 2.723E05 | 2.133E-53 |
| H | 3 | gzip ICR | Levene | 1.077E01 | 1.060E-03 |
| H | 3 | H.264 ICR | Mann-Whitney U | 2.263E05 | 1.484E-15 |
| H | 3 | H.264 ICR | Levene | 1.886E01 | 1.523E-05 |
| H | 3 | MPEG-2 ICR | Mann-Whitney U | 2.624E05 | 6.028E-43 |
| H | 3 | MPEG-2 ICR | Levene | 6.819E01 | 3.880E-16 |
| H | 3 | Firing Rates | Mann-Whitney U | 2.780E05 | 6.093E-60 |
| H | 3 | Firing Rates | Levene | 1.213E02 | 6.004E-27 |
| H | 3 | Fano Factor | Mann-Whitney U | 2.576E05 | 2.908E-38 |
| H | 3 | Fano Factor | Levene | 1.874E00 | 1.713E-01 |
| H | 3 | PCA | Mann-Whitney U | 3.254E04 | 2.929E-133 |
| H | 3 | PCA | Levene | 1.632E02 | 3.902E-35 |
| H | 3 | FA | Mann-Whitney U | 4.121E03 | 1.011E-188 |
| H | 3 | FA | Levene | 3.015E-03 | 9.562E-01 |

## Notes

### Competing Interest Statement

The authors have declared no competing interest.

## References

[1] I. H. Stevenson and K. P. Kording, “How advances in neural recording affect data analysis,” Nature Neuroscience, vol. 14, p. 139–142, Jan. 2011.

[2] J. P. Cunningham and B. M. Yu, “Dimensionality reduction for large-scale neural recordings,” Nature Neuroscience, vol. 17, p. 1500–1509, Aug. 2014.

[3] S. Vyas, M. D. Golub, D. Sussillo, and K. V. Shenoy, “Computation through neural population dynamics,” Annual Review of Neuroscience, vol. 43, p. 249–275, July 2020.

[4] R. B. Ebitz and B. Y. Hayden, “The population doctrine in cognitive neuroscience,” Neuron, vol. 109, p. 3055–3068, Oct. 2021.

[5] P. Gao, E. Trautmann, B. Yu, G. Santhanam, S. Ryu, K. Shenoy, and S. Ganguli, “A theory of multineuronal dimensionality, dynamics and measurement,” Nov. 2017.

[6] F. Pei, J. Ye, D. Zoltowski, A. Wu, R. H. Chowdhury, H. Sohn, J. E. O’Doherty, K. V. Shenoy, M. T. Kaufman, M. Churchland, M. Jazayeri, L. E. Miller, J. Pillow, I. M. Park, E. L. Dyer, and C. Pandarinath, “Neural latents benchmark ‘21: Evaluating latent variable models of neural population activity,” 2021.

[7] J. C. Kao, P. Nuyujukian, S. I. Ryu, M. M. Churchland, J. P. Cunningham, and K. V. Shenoy, “Single-trial dynamics of motor cortex and their applications to brain-machine interfaces,” Nature Communications, vol. 6, July 2015.

[8] M. R. Keshtkaran, A. R. Sedler, R. H. Chowdhury, R. Tandon, D. Basrai, S. L. Nguyen, H. Sohn, M. Jazayeri, L. E. Miller, and C. Pandarinath, “A large-scale neural network training framework for generalized estimation of single-trial population dynamics,” Nature Methods, vol. 19, p. 1572–1577, Nov. 2022.

[9] C. E. Shannon, “A mathematical theory of communication,” Bell System Technical Journal, vol. 27, p. 379–423, July 1948.

[10] T. M. Cover and J. A. Thomas, Elements of Information Theory. Wiley, Apr. 2005.

[11] M. Hayashi, Quantum Information Theory: Mathematical Foundation. Springer Berlin Heidelberg, 2017.

[12] T. D. Schneider, “A brief review of molecular information theory,” Nano Communication Networks, vol. 1, p. 173–180, Sept. 2010.

[13] A. Borst and F. E. Theunissen, “Information theory and neural coding,” Nature Neuroscience, vol. 2, p. 947–957, Nov. 1999.

[14] D. A. Butts, C. Weng, J. Jin, C.-I. Yeh, N. A. Lesica, J.-M. Alonso, and G. B. Stanley, “Temporal precision in the neural code and the timescales of natural vision,” Nature, vol. 449, p. 92–95, Sept. 2007.

[15] S. E. Clarke, A. Longtin, and L. Maler, “The neural dynamics of sensory focus,” Nature Communications, vol. 6, Nov. 2015.

[16] P. Wollstadt, K. K. Sellers, L. Rudelt, V. Priesemann, A. Hutt, F. Fröhlich, and M. Wibral, “Breakdown of local information processing may underlie isoflurane anesthesia effects,” PLOS Computational Biology, vol. 13, p. e1005511, June 2017.

[17] M. Hernaez, D. Pavlichin, T. Weissman, and I. Ochoa, “Genomic data compression,” Annual Review of Biomedical Data Science, vol. 2, p. 19–37, July 2019.

[18] B. J. Kagan, A. C. Kitchen, N. T. Tran, F. Habibollahi, M. Khajehnejad, B. J. Parker, A. Bhat, B. Rollo, A. Razi, and K. J. Friston, “In vitro neurons learn and exhibit sentience when embodied in a simulated game-world,” Neuron, vol. 110, pp. 3952–3969.e8, Dec. 2022.

[19] N. M. Timme and C. Lapish, “A tutorial for information theory in neuroscience,” eneuro, vol. 5, pp. ENEURO.0052–18.2018, May 2018.

[20] E. D. Fagerholm, Z. Dezhina, R. J. Moran, F. E. Turkheimer, and R. Leech, “A primer on entropy in neuroscience,” Neuroscience amp; Biobehavioral Reviews, vol. 146, p. 105070, Mar. 2023.

[21] A. Luczak, “Entropy of neuronal spike patterns,” Entropy, vol. 26, p. 967, Nov. 2024.

[22] S. Kulkarni and D. S. Bassett, “Toward principles of brain network organization and function,” Annual Review of Biophysics, vol. 54, p. 353–378, May 2025.

[23] L. Yamada, T. Oskotsky, and P. Nuyujukian, “Compression-enabled joint entropy estimation for seizure detection on human intracortical electroencephalography,” IEEE Transactions on Biomedical Engineering, pp. 3440–3452, Dec 2015.

[24] D. Salomon, A Concise Introduction to Data Compression. Springer London, 2008.

[25] M. Nelson and J.-L. Gailly, The data compression book (2nd ed.). USA: MIS:Press, 1995.

[26] I. E. G. Richardson, “H.264 and mpeg-4 video compression: Video coding for next generation multimedia,” 2003.

[27] U. Bihr, H. Xu, C. Bulach, M. Lorenz, J. Anders, and M. Ortmanns, “Real-time data compression of neural spikes,” in 2014 IEEE 12th International New Circuits and Systems Conference (NEWCAS), p. 436–439, IEEE, June 2014.

[28] M. Pagin and M. Ortmanns, “A neural data lossless compression scheme based on spatial and temporal prediction,” in 2017 IEEE Biomedical Circuits and Systems Conference (BioCAS), p. 1–4, IEEE, Oct. 2017.

[29] P. Yan, A. Akhoundi, N. P. Shah, P. Tandon, D. G. Muratore, E. J. Chichilnisky, and B. Murmann, “Data compression versus signal fidelity tradeoff in wired-or analog-to-digital compressive arrays for neural recording,” IEEE Transactions on Biomedical Circuits and Systems, vol. 17, p. 754–767, Aug. 2023.

[30] J. M. Amigó, J. Szczepański, E. Wajnryb, and M. V. Sanchez-Vives, “Estimating the entropy rate of spike trains via lempel-ziv complexity,” Neural Computation, vol. 16, p. 717–736, Apr. 2004.

[31] D. Abásolo, S. Simons, R. Morgado da Silva, G. Tononi, and V. V. Vyazovskiy, “Lempel-ziv complexity of cortical activity during sleep and waking in rats,” Journal of Neurophysiology, vol. 113, p. 2742–2752, Apr. 2015.

[32] M. M. Schartner, R. L. Carhart-Harris, A. B. Barrett, A. K. Seth, and S. D. Muthukumaraswamy, “Increased spontaneous meg signal diversity for psychoactive doses of ketamine, lsd and psilocybin,” Scientific Reports, vol. 7, Apr. 2017.

[33] J.-P. Noel, Y. Ishizawa, S. R. Patel, E. N. Eskandar, and M. T. Wallace, “Leveraging nonhuman primate multisensory neurons and circuits in assessing consciousness theory,” The Journal of Neuroscience, vol. 39, p. 7485–7500, July 2019.

[34] A. G. Casali, O. Gosseries, M. Rosanova, M. Boly, S. Sarasso, K. R. Casali, S. Casarotto, M.-A. Bruno, S. Laureys, G. Tononi, and M. Massimini, “A theoretically based index of consciousness independent of sensory processing and behavior,” Science Translational Medicine, vol. 5, Aug. 2013.

[35] R. Comolatti, A. Pigorini, S. Casarotto, M. Fecchio, G. Faria, S. Sarasso, M. Rosanova, O. Gosseries, M. Boly, O. Bodart, D. Ledoux, J.-F. Brichant, L. Nobili, S. Laureys, G. Tononi, M. Massimini, and A. G. Casali, “A fast and general method to empirically estimate the complexity of brain responses to transcranial and intracranial stimulations,” Brain Stimulation, vol. 12, p. 1280–1289, Sept. 2019.

[36] M. Rosanova, S. Casarotto, C. Derchi, G. Hassan, S. Russo, S. Sarasso, A. Vigano, M. Massimini, and A. Comanducci, “The perturbational complexity index detects capacity for consciousness earlier than the recovery of behavioral responsiveness in subacute brain-injured patients,” Brain Stimulation, vol. 16, p. 371, Jan. 2023.

[37] M. London, A. Schreibman, M. Häusser, M. E. Larkum, and I. Segev, “The information efficacy of a synapse,” Nature Neuroscience, vol. 5, p. 332–340, Mar. 2002.

[38] Y. Tang and D. Durand, “A tunable support vector machine assembly classifier for epileptic seizure detection,” Expert Systems with Applications, vol. 39, p. 3925–3938, Mar. 2012.

[39] M. Zbili and S. Rama, “A quick and easy way to estimate entropy and mutual information for neuroscience,” Frontiers in Neuroinformatics, vol. 15, June 2021.

[40] P. N. W. G. Deutsch, “GZIP file format specification version 4.3,” 1996.

[41] P. Tudor, “Mpeg-2 video compression tutorial,” in IEE Colloquium on MPEG-2 - What it is and What it isn’t, vol. 1995, p. 2–2, IEE, 1995.

[42] D. Marpe, T. Wiegand, and G. Sullivan, “The h.264/mpeg4 advanced video coding standard and its applications,” IEEE Communications Magazine, vol. 44, no. 8, pp. 134–143, 2006.

[43] J. P. Cunningham, V. Gilja, S. I. Ryu, and K. V. Shenoy, “Methods for estimating neural firing rates, and their application to brain–machine interfaces,” Neural Networks, vol. 22, p. 1235–1246, Nov. 2009.

[44] K. Rajdl, P. Lansky, and L. Kostal, “Fano factor: A potentially useful information,” Frontiers in Computational Neuroscience, vol. 14, Nov. 2020.

[45] A. Georgopoulos, J. Kalaska, R. Caminiti, and J. Massey, “On the relations between the direction of two-dimensional arm movements and cell discharge in primate motor cortex,” The Journal of Neuroscience, vol. 2, p. 1527–1537, Nov. 1982.

[46] A. B. Schwartz, “Useful signals from motor cortex,” The Journal of Physiology, vol. 579, p. 581–601, Mar. 2007.

[47] P. J. Marino, L. Bahureksa, C. F. Fisac, E. R. Oby, A. L. Smoulder, A. Motiwala, A. D. Degenhart, E. M. Grigsby, W. M. Joiner, S. M. Chase, B. M. Yu, and A. P. Batista, “A posture subspace in the primary motor cortex,” Neuron, vol. 113, pp. 3647–3660.e10, Nov. 2025.

[48] I. E. Bray, S. E. Clarke, K. M. Casey, and P. Nuyujukian, “Neuroelectrophysiology-compatible electrolytic lesioning,” eLife, vol. 12, July 2024.

[49] D. A. Adamos, E. K. Kosmidis, and G. Theophilidis, “Performance evaluation of pca-based spike sorting algorithms,” Computer Methods and Programs in Biomedicine, vol. 91, p. 232–244, Sept. 2008.

[50] P. N. W. G. Deutsch, “DEFLATE Compressed Data Format Specification version 1.3,” 1996.

[51] J. Ziv and A. Lempel, “A universal algorithm for sequential data compression,” IEEE Transactions on Information Theory, vol. 23, p. 337–343, May 1977.

[52] D. Huffman, “A method for the construction of minimum-redundancy codes,” Proceedings of the IRE, vol. 40, p. 1098–1101, Sept. 1952.

[53] S. R. Ely, “Mpeg video coding: A simple introduction,” EBU Technical Review, 1995.

[54] H. H. Barbero, M. and N. D. Wells, “Dct source coding and current implementations for hdtv,” EBU Technical Review, 1992.

[55] B. G. Haskell, P. G. Howard, Y. A. LeCun, A. Puri, J. Ostermann, M. Civanlar, L. Rabiner, L. Bottou, and P. Haffner, “Image and video coding—emerging standards and beyond,” in Readings in Multimedia Computing and Networking (K. Jeffay and H. Zhang, eds.), The Morgan Kaufmann Series in Multimedia Information and Systems, pp. 89–112, San Francisco: Morgan Kaufmann, 2002.

[56] H. Wang, S. Kwong, and C.-W. Kok, “Efficient prediction algorithm of integer dct coefficients for h.264/avc optimization,” IEEE Transactions on Circuits and Systems for Video Technology, vol. 16, p. 547–552, Apr. 2006.

[57] A. Samcovic and J. Turan, “H.264/avc multiple reference video frames,” in 2008 International Conference on Signals and Electronic Systems, p. 431–434, IEEE, 2008.

[58] N. S. Narayanan, E. Y. Kimchi, and M. Laubach, “Redundancy and synergy of neuronal ensembles in motor cortex,” The Journal of Neuroscience, vol. 25, p. 4207–4216, Apr. 2005.

[59] M. T. Kaufman, M. M. Churchland, S. I. Ryu, and K. V. Shenoy, “Cortical activity in the null space: permitting preparation without movement,” Nature Neuroscience, vol. 17, p. 440–448, Feb. 2014.

[60] L. N. Driscoll, N. L. Pettit, M. Minderer, S. N. Chettih, and C. D. Harvey, “Dynamic reorganization of neuronal activity patterns in parietal cortex,” Cell, vol. 170, pp. 986–999.e16, Aug. 2017.

[61] J. A. Hennig, M. D. Golub, P. J. Lund, P. T. Sadtler, E. R. Oby, K. M. Quick, S. I. Ryu, E. C. Tyler-Kabara, A. P. Batista, B. M. Yu, and S. M. Chase, “Constraints on neural redundancy,” eLife, vol. 7, Aug. 2018.

[62] B. Campos, H. Choi, A. T. DeMarco, A. Seydell-Greenwald, S. J. Hussain, M. T. Joy, P. E. Turkeltaub, and W. Zeiger, “Rethinking remapping: Circuit mechanisms of recovery after stroke,” The Journal of Neuroscience, vol. 43, p. 7489–7500, Nov. 2023.

[63] M. D. Golub, P. T. Sadtler, E. R. Oby, K. M. Quick, S. I. Ryu, E. C. Tyler-Kabara, A. P. Batista, S. M. Chase, and B. M. Yu, “Learning by neural reassociation,” Nature Neuroscience, vol. 21, p. 607–616, Mar. 2018.

[64] R. J. Nudo, “Mechanisms for recovery of motor function following cortical damage,” Current Opinion in Neurobiology, vol. 16, p. 638–644, Dec. 2006.

[65] M. T. Joy and S. T. Carmichael, “Encouraging an excitable brain state: mechanisms of brain repair in stroke,” Nature Reviews Neuroscience, vol. 22, p. 38–53, Nov. 2020.

[66] N. Mohsenian, R. Rajagopalan, and C. A. Gonzales, “Single-pass constant- and variable-bit-rate mpeg-2 video compression,” IBM Journal of Research and Development, vol. 43, p. 489–509, July 1999.

[67] G. W. Fraser, S. M. Chase, A. Whitford, and A. B. Schwartz, “Control of a brain–computer interface without spike sorting,” Journal of Neural Engineering, vol. 6, p. 055004, Sept. 2009.

[68] C. A. Chestek, V. Gilja, P. Nuyujukian, J. D. Foster, J. M. Fan, M. T. Kaufman, M. M. Churchland, Z. Rivera-Alvidrez, J. P. Cunningham, S. I. Ryu, and K. V. Shenoy, “Long-term stability of neural prosthetic control signals from silicon cortical arrays in rhesus macaque motor cortex,” Journal of Neural Engineering, vol. 8, p. 045005, July 2011.

[69] B. P. Christie, D. M. Tat, Z. T. Irwin, V. Gilja, P. Nuyujukian, J. D. Foster, S. I. Ryu, K. V. Shenoy, D. E. Thompson, and C. A. Chestek, “Comparison of spike sorting and thresholding of voltage waveforms for intracortical brain–machine interface performance,” Journal of Neural Engineering, vol. 12, p. 016009, Dec. 2014.

[70] E. M. Trautmann, S. D. Stavisky, S. Lahiri, K. C. Ames, M. T. Kaufman, D. J. O’Shea, S. Vyas, X. Sun, S. I. Ryu, S. Ganguli, and K. V. Shenoy, “Accurate estimation of neural population dynamics without spike sorting,” Neuron, vol. 103, pp. 292–308.e4, July 2019.

[71] J. Dai, P. Zhang, H. Sun, X. Qiao, Y. Zhao, J. Ma, S. Li, J. Zhou, and C. Wang, “Reliability of motor and sensory neural decoding by threshold crossings for intracortical brain–machine interface,” Journal of Neural Engineering, vol. 16, p. 036011, Apr. 2019.

[72] N. Sagan and T. Weissman, “A family of lz78-based universal sequential probability assignments,” 2024.

[73] H. Inan, M. A. Erdogdu, and M. Schnitzer, “Robust estimation of neural signals in calcium imaging,” in Advances in Neural Information Processing Systems (I. Guyon, U. V. Luxburg, S. Bengio, H. Wallach, R. Fergus, S. Vishwanathan, and R. Garnett, eds.), vol. 30, Curran Associates, Inc., 2017.

[74] K. H. Lee, Y.-L. Ni, J. Colonell, B. Karsh, J. Putzeys, M. Pachitariu, T. D. Harris, and M. Meister, “Electrode pooling can boost the yield of extracellular recordings with switchable silicon probes,” Nature Communications, vol. 12, Sept. 2021.

